# Classification of motor-related brain states using high frequency information from muscle recordings

**DOI:** 10.1101/2025.03.11.642676

**Authors:** B Zicher, J Ibáñez, D Farina

**Affiliations:** Department of Computing and Bioengineering, Imperial College London, London, UK; BSICoS group, I3A Institute, IIS Aragón, University of Zaragoza, Zaragoza, Spain; Centro de Investigación Biomédica en Red en Bioingeniería, Biomateriales y Nanomedicina (CIBER-BBN), Zaragoza, Spain

## Abstract

Motor neural interfaces use recordings from the nervous system to extract control signals used to interact with the environment. Muscle signals are becoming an increasingly popular choice for motor interfaces, especially when used to estimate the neural motor commands driving muscle contractions and eventually movement. Here we study the possibility of using electromyography (EMG) to classify motor-related cortical states associated with the cancellation of movement in a ‘GO’/’NO-GO’ task, which has been previously linked to changes in cortical beta oscillations. We show that beta (13-30Hz) and low gamma (30-45Hz) frequency bands have the most predictive power in electroencephalography (EEG) and EMG recordings. Moreover, we found comparable accuracy in cancellation state decoding from EEG (average of 74%) and EMG (average of 77%) recordings, which supports the concept of using peripheral signals to predict cortical activity associated to specific motor-related brain states.

## 1 Introduction

Cortical oscillatory activity in beta (13-30Hz) band has been studied in motor tasks such as steady contractions, movement preparation, cancellation and motor imagery (Barone and Rossiter, 2021; Engel and Fries, 2010; Kilavik et al., 2013). Such oscillatory activity may be recorded by non-invasive brain-machine interfaces (BMIs) and used in neurofeedback paradigms (Abiri et al., 2019; Cook et al., 2021; Wolpaw et al., 2002). However, non-invasive brain recordings of beta activity vary across subjects and are susceptible to background noise, which limits their real-world applicability (Martini et al., 2020).

It has been shown that cortical beta and gamma (>30Hz) frequencies are reliably and partly linearly transmitted to the peripheral nervous system. This is mainly supported by the observed high levels of coherence between brain and muscle recordings in these frequency bands in both humans and primates (Baker, 2007; Conway et al., 1995; Schoffelen et al., 2005, 2011). Even though it is well known that spinal motor neurons receive inputs in frequencies outside the bandwidth for muscle control (>10 Hz), there have been little efforts in decoding high-frequency inputs and using them for interfacing purposes (Farina et al., 2014). Recent work points towards the prospect of extracting beta band oscillations in real-time from motor unit (MU) population activity (Bräcklein et al., 2021; Bräcklein et al., 2022).

Here we wanted to investigate whether muscle signals can be used to distinguish on a trial-by-trial basis different motor-related cortical states associated with changes in the power of neural rhythms. For this, we used previously acquired data from healthy subjects during a ‘GO’/’NO-GO’ paradigm. The task required subjects to prepare and then produce or cancel a motor command with their tibialis anterior (TA) muscle, all while holding a stable contraction. It has been shown that the state of cancellation of a prepared movement can cause large changes in the power of brain rhythms that are in turn reflected in the oscillatory activity recorded in the muscles (Zicher et al., 2024). However, it is unclear how consistent the changes are on a trial-by-trial basis and whether they are salient enough to classify a state based on surface EMG (sEMG) recordings. We defined two states: *baseline*, when subjects were holding a stable contraction, and *cancellation*, representing the post ‘NO-GO’ period in which subjects had to maintain the sustained contraction of the TA unaltered. We found that the classification accuracy in distinguishing between these two states is comparable when using EMG or EEG. Moreover, we further confirmed that the high frequencies are most relevant for distinguishing the *baseline* from *cancellation*.

These results are a proof-of-concept for the potential use of EMG-based peripheral interfaces in extracting and classifying motor-related brain states.

## 2 Materials and methods

### 2.1 Dataset

The dataset used in this work has been previously published in Zicher et al., 2024. The procedures will be briefly explained in the following.

#### Ethics

Data from 7 participants (all male; ages 21-38 yrs) was collected. The participants gave a written informed consent to take part in the experiments. The study was approved by the Imperial College Ethics committee (18IC4685) and was done in accordance with the Declaration of Helsinki.

#### Data collection

Electroencephalography (EEG), high-density surface electromyography (HD-EMG) and force data were acquired concurrently during an ankle dorsiflexion task.

EEG signals were recorded from 31 scalp positions using active gel-based electrodes. The electrodes were arranged in accordance with the International 10–20 system and FCz was used as the reference (actiCAP, Brain Products GmbH). The signals were amplified and sampled at 1kHz using BrainVision actiCHamp Plus (Brain Products GmbH).

EMG signals were recorded from the tibialis anterior of the right leg using four 64-channel surface grids (13×5 arrangement, 4-mm interelectrode distance; manufactured by OT Bioelettronica, IT), placed in two columns to cover the area of the muscle. The recorded signals were amplified and sampled at 2048Hz using a Quattrocento amplifier (OT Bioelettronica, IT). Together with the EMG signals, ankle dorsiflexion forces measured with an ankle dynamo-meter (TF-022, CCT Transducer) sampled at 1kHz. Throughout the experiment, subjects sat on a chair with their right knee at 75°, foot fixed to an ankle dynamometer, which had the force transducer on the pedal.

Subjects received visual feedback about their force levels displayed on a monitor placed in front of them.

The recorded signals were synchronised using a common digital trigger.

#### Task

At the beginning of the recording session, the maximum voluntary contraction (MVC) force that each subject could produce doing a dorsiflexion of the ankle was measured. Force feedback was given in %MVC in the main experimental parts following this first step. The main part of the study consisted of three identical blocks with 35 trials each. Each trial could be either a ‘GO’ or a ‘NO-GO’ trial with equal probability. Participants were guided throughout the trials by auditory cues of various frequencies and durations. The first cue, at the beginning of the trial, indicated to subjects that they had to produce a 10% MVC isometric contraction. The second cue (warning stimulus) announced that the imperative stimulus was coming and subjects had to prepare a ballistic contraction. One second after hearing the third cue, participants either had to keep the stable contraction (’NO-GO’ trials), or to go ahead with the prepared action (’GO’ trials) (Figure 1). Finally, a last cue indicated the end of a trial, when subjects had to go back to a relaxed position until the next trial. Participants were trained on the task until they were confident in their abilities.

**Figure 1:**
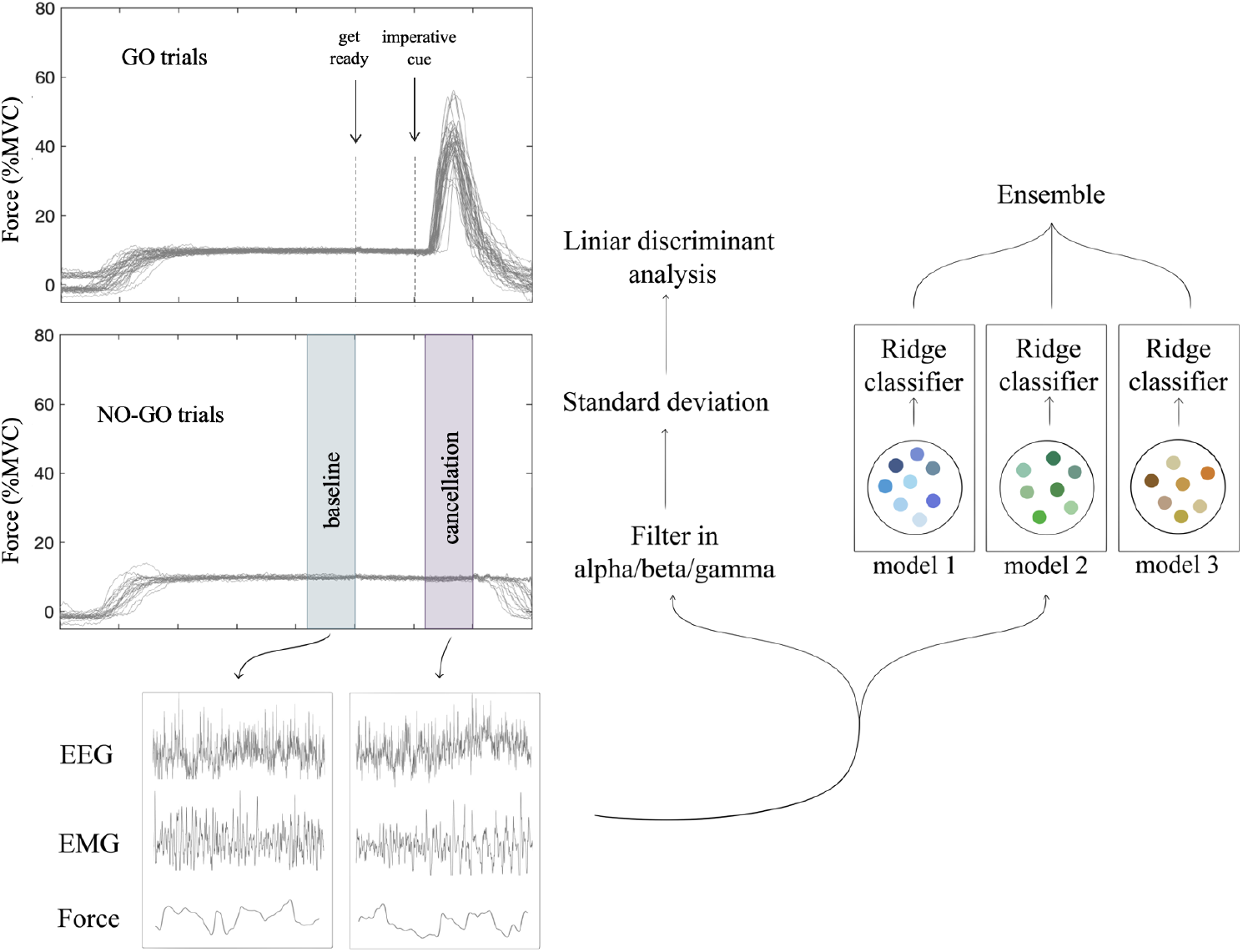
Schematic of the task considered in the present study and the overall data processing methodology. Subjects were asked to react to auditory signals while holding a stable contraction with their tibialis anterior muscle. In the ‘GO’ trials, they produced a ballistic contraction when hearing the imperative cue, while in ‘NO-GO’ trials they were instructed to keep the force stable. Force, EEG and EMG signals were extracted from *baseline* and *cancellation* periods. For the first set of analyses, Linear Discriminant Analysis (LDA) models were trained on the feature extracted as standard deviation of the filtered signal from specific frequency bands. In the second set of analyses, an ensemble of models was created, with each type of model extracting different features from the data and then training a ridge classifier. Only one of each of the three feature extraction types are shown in the figure for simplicity.

#### Signal processing

We tested the classification of *baseline* and *cancellation* states from EEG, EMG and force recordings. The *baseline* state represents the period where participants were only holding a stable contraction, while *cancellation* is defined as the period right after the ‘NO-GO’ cue, when the second prepared action is aborted. Data segments −1.8s to −1s (*baseline*) and 0.2s to 1s (*cancellation*), relative to the imperative cue, were extracted from brain, muscle and force recordings (Figure 1). These windows were chosen since the average reaction time for the ‘GO’ trials is above 250ms (Zicher et al., 2024), thus we were not expecting changes earlier than 200ms post-cue. Only valid trials were used (see Zicher et al., 2024).

In order to keep results comparable between recording types, we created one channel per modality. For a global analysis of EMG, we obtained a single signal by averaging all EMG channels. For EEG, the Laplacian derivation from channel ‘Cz’ was computed by subtracting the average electric potential recorded from the four closest equidistant electrodes (i.e., FC1, FC2, CP1, CP2) (Kayser and Tenke, 2015). This channel was chosen as it produced the largest cortico-muscular coherence values in high frequencies. Features from this new channel were used to train models. The goal of this work was not to quantify the best accuracy that could be achieved, but to evaluate whether better than chance classification is possible using EMG signals.

For the classification based on information from different bandwidths, EEG, EMG and force signals were first band-pass filtered using a 3rd order Butterworth zero-phase filter in the alpha (10-13 Hz), beta (13-30 Hz) and gamma (30-45 Hz) bands. The filtered signals were then segmented to extract data from the *baseline* and *cancellation* periods according to the above-mentioned intervals, and the standard deviation (SD) of the extracted signals was calculated for each trial. This SD value was then used as input to the classification model used.

### 2.2 Algorithms

Two sets of analyses were performed. The first classified states based on the strength of the signals in different frequency bands; the second used the raw recorded signals directly for feature extraction by the model used for classification (EMG was recorded as high-pass filtered at >10Hz). Classification algorithms and optimisations were implemented in Python. Subject-specific models were trained.

A Linear Discriminant Analysis (LDA) classifier with automatic shrinkage with Ledoit-Wolf lemma was used in the first part of this study. The feature used as input to these models was the standard deviation of the filtered signal (see Signal processing above).

An ensemble of 12 classifiers was created with three types of transforms designed for time series data for the second set of analyses. The first type used a RandOm Convolutional KErnel Transform (ROCKET), the second one, a MINImally RandOm Convolutional KErnel Transform (MiniRocket), and the third one, a Word ExtrAction for time SEries cLassification (WEASEL) 2.0 Transform (Dempster et al., 2020, 2021; Schäfer and Leser, 2023). WEASEL 2.0 is based on the Symbolic Fourier Approximation (SFA) Transform. The features extracted with these transformers were scaled and used by a ridge classifier to predict the state. Four classifiers were trained for each type of transform, with the regularization strength of the classifiers chosen through cross-validation. During predictions, each classifiers prediction was weighted by the accuracy achieved during cross-validation, as done previously for an ensemble of ROCKET classifiers (Middlehurst et al., 2021).

### 2.3 Optimisation

For each of the classification accuracy results, stratified 5-fold cross-validation was run 30 times and results averaged. This was done to get a robust estimate of the average accuracy per subject.

#### Controls

Given the small dataset, randomness can have a large effect on the accuracy achieved on a data distribution. Therefore, we quantified the distribution of accuracy that was achieved with random labels to estimate the chance level. For this purpose, class labels were randomized (total of 50 times), the 5-fold cross-validation was run 30 times and results averaged. This provided a distribution of accuracy for the control case (chance-level predictions). When models were trained on the real labels, the classification was considered better than chance level if the accuracy was above the upper bound of the 95% confidence interval. This was checked by testing if 97.5% of control accuracy values were smaller than the classification accuracy on real labels.

## 3 Results

For the subject-specific models, on average 41 (±6) trials were included in the analysis.

### 3.1 Analysis 1

With the first analysis we evaluated the predictive power of alpha, beta and gamma frequencies in EEG, EMG and force recordings. For this purpose, LDA classifiers were trained on the SD of signals in the various bandwidths.

The results from the subject-specific models suggest that contents in the beta and gamma bands in the EEG and EMG signals have the highest predictive power (Figure 2). Only in one out of the seven subjects EMG did not result in classifications above chance level. Accuracies were highly variable across subjects, with values ranging from 58% to 91%.

**Figure 2:**
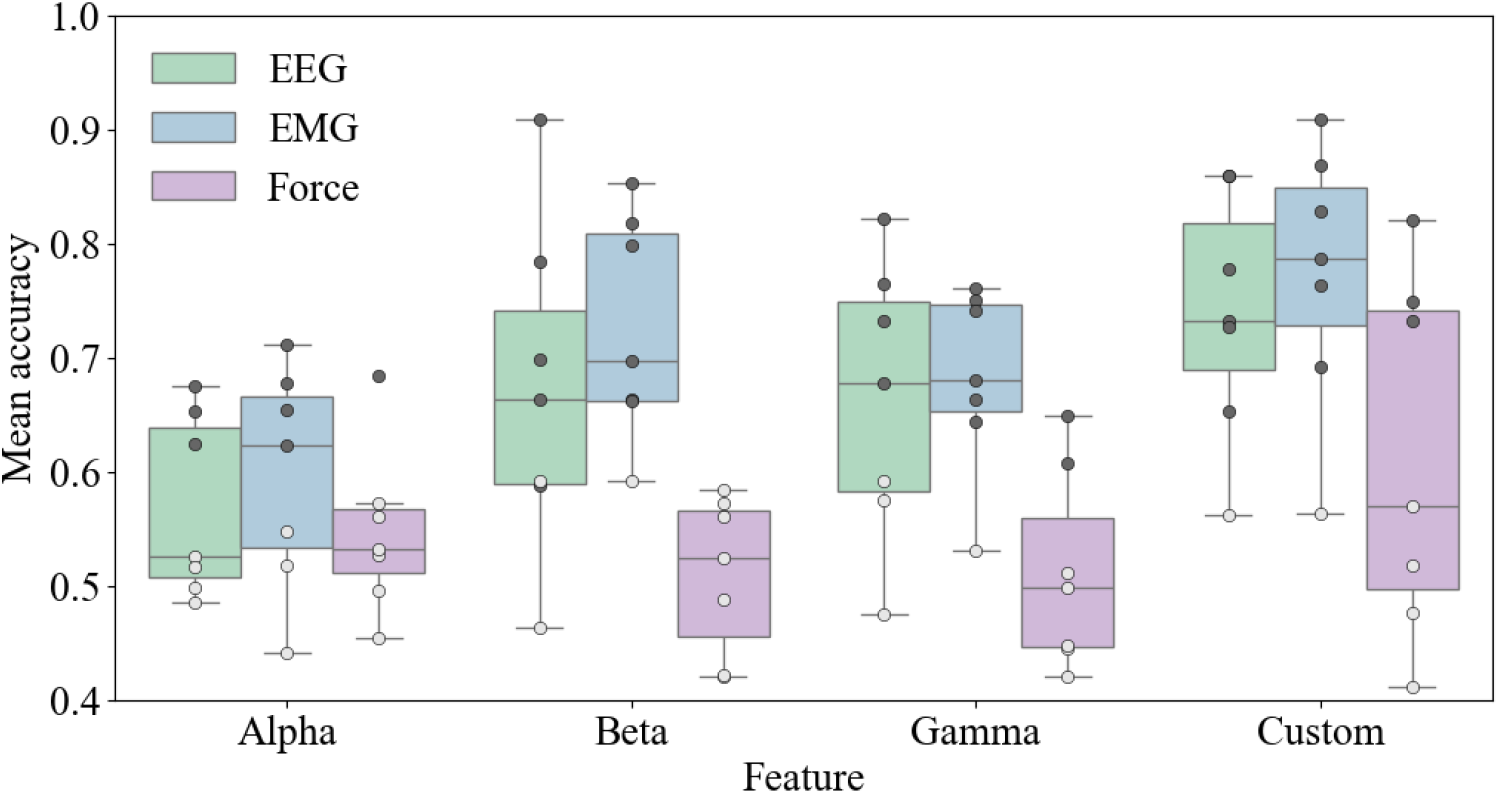
Test accuracies of models trained and tested on individual subject data. Results are grouped by the features used by the models. Each point represents a subject. Accuracy values not above chance level are plotted in white, while those significant are plotted in dark grey.

### 3.2 Analysis 2

In the next analysis, we used more complex models made for time series classification to study if accuracies can be improved when features are not restricted to specific bandwidths. An ensemble of models was trained for each subject separately. On average, the ensemble achieved a higher accuracy for both EEG and EMG data, compared to the LDA classifier trained on beta feature from Analysis 1 (Figure 2). The mean accuracy increased from 67% to 74% for EEG and from 73% to 77% for EMG. The results from classifying with EEG and EMG were comparable, with EMG providing higher accuracies in four out of seven subjects. For some subjects, force was also predictive of state, but overall the results were notably worse than with EMG or EEG features. Moreover, the accuracy when classifying states by using the custom force features was not significantly correlated to the accuracy achieved when using custom EMG features, thus possible small force changes across states did not fully explain the results (Figure 3B). The correlation between accuracies achieved with beta EMG and custom force features classification was not significant either (p=0.879; not shown in figure). Conversely, the accuracy obtained from only the beta band EMG classification was correlated to the accuracy achieved from classifying with custom EMG features (Figure 3A), indicating that the main discriminative information was in the beta band.

**Figure 3:**
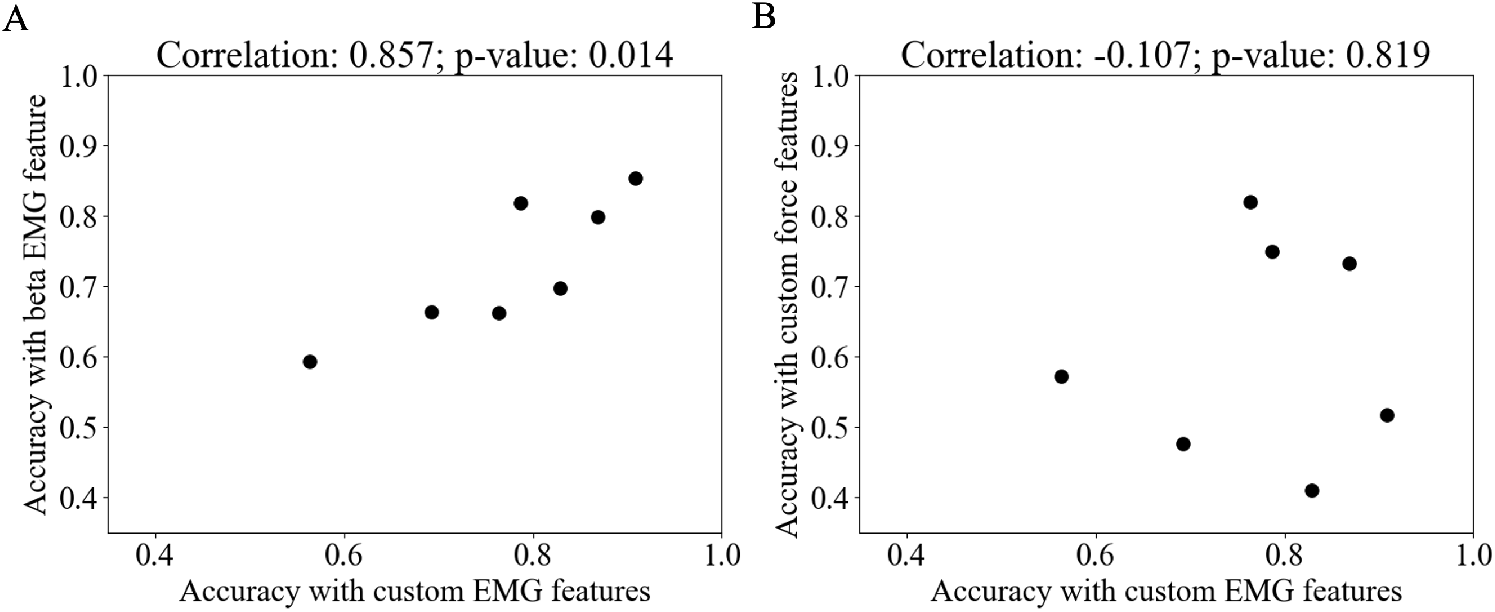
Spearman correlation between test accuracy when using different signals and features. Panel A shows the accuracy achieved by individual subject models trained with custom EMG features against the models trained with the beta feature. Panel B shows results for the models trained with EMG vs force features. Each point represents result for one participant.

## 4 Discussion

In this study we assessed if it was possible to distinguish two motor-related states known to have different oscillatory activity using muscle signals. This was motivated by recent studies showing that high-frequency rhythms (in the beta and gamma range) in the motor cortex can be transmitted to muscles in the periphery in a linear and robust way (Ibáñez et al., 2024; Zicher et al., 2024).

Based on this, the study tested for the first time if muscle recordings (EMG) could be used to detect brain oscillations and classify motor-related cortical states. Specifically, we aimed at distinguishing a state of stable muscle contraction (*baseline*) from the state of preparing and aborting a second movement on top of the isometric contraction (*cancellation*). Our results show that EMG-based classification of the studied states led to accuracies that were comparable to those obtained using signals directly recorded from the brain using EEG. These results represent a proof-of-concept for a peripheral interface driven by cortical oscillations.

The first analysis evaluated the predictive power of the activity in the alpha, beta and gamma bands in the EEG, EMG and force measurements. We found that beta and gamma frequencies measured with EEG and EMG had significant predictive power regarding the difference between the two states classified. Previous work has shown that, on average, there are increases in both beta and gamma frequencies during the period after a ‘NO-GO’ instruction in a warned ‘GO’/’NO-GO’ task (Zicher et al., 2024). As a continuation of that work, here we show that it is possible to use cortical or peripheral recordings to classify these two states on a trial-by-trial basis. In the context of physiological signals, classification with the beta and gamma band activity in EEG and EMG was comparable, which suggests that the information captured by brain and muscle recordings are similar. Given that the results presented here were achieved by averaging EMG channels, future work should further explore what is the best minimal or optimal electrode setup that could be used to reliably classify motor-related states based on the oscillations transmitted to the muscles. Furthermore, the potential advantages of multimodal (brain and muscle) models should be explored. In addition, for larger datasets, decomposing EMG into constituent spike trains and using that as input to models could have an advantage over EMG, which is highly subject-specific (Kapelner et al., 2019).

In order not to restrict the features to specific frequencies and test how much additional information the full bandwidth provides, in the second analysis of this work we used a custom ensemble of kernel and dictionary-based models. Using this approach, we achieved a slight increase in accuracy for all recording types. Predictions based on EMG information compared to EEG, were better in four out of seven subjects. The correlation of accuracy between individual subject models trained on beta from EMG and full bandwidth EMG was 0.857, indicating the importance of beta power changes when classifying the state of *cancellation* (Figure 3).

In the case of models trained with force data, one and two out of seven, for alpha and gamma frequencies respectively, had accuracies greater than chance level. The participant with changes in alpha frequencies also had changes in gamma. Given the complexity of the task, it was impossible to completely restrict the force output of the participants. However, the changes in force did not seem to reflect the changes in high frequencies seen in the EMG. Indeed, for one of the participants, force was slightly and steadily increasing throughout the *cancellation* window, something not seen in other participants and probably not related to the *cancellation* state, but likely picked up by the ensemble model. Moreover, there was no significant correlation between classification accuracies with EMG and force, suggesting that changes in force (on an individual subject level) are not correlated with the changes found in EMG during the *cancellation* state.

It is worth noting that even though only the correct trials (where participants reacted correctly to cues) were kept when labeling data, it was assumed that subjects were indeed in the correct state. However, it is hard to quantify how engaged the subjects were with the task on a trial-by-trial basis. No change in force produced in the ‘NO-GO’ trials does not necessarily mean that participants prepared and canceled a second movement. It is unclear how differences in focus would affect the saliency of changes we found between the two states and whether it contributed to inter-subject variability. Furthermore, it raises the question whether the magnitude and consistency of these modulations is fixed or can be trained and improved.

## Limitations

Classification was tested for seven participants, with a relatively small number of trials. While accuracies (with EEG and EMG) were consistently better than chance, the goal of study was not to quantify the best possible accuracy that can be achieved with these two recording types. Future work should extend the findings of this work to larger participant groups. It is likely that with a bigger dataset, the classification accuracies would improve, with the possibility of using deep learning approaches.

When considering a real-world application of a peripheral interface, the classification of a self-paced state may be preferred. In this study, the *cancellation* state followed an external cue, thus future research should explore other tasks, where participants are not restricted by cues.

